# Embryo CHH hypermethylation is mediated by RdDM and is autonomously directed in *Brassica rapa*

**DOI:** 10.1101/2020.08.26.268573

**Authors:** Tania Chakraborty, Timmy Kendall, Jeffrey W. Grover, Rebecca A. Mosher

**Affiliations:** The School of Plant Sciences, The University of Arizona, Tucson, AZ 85721 USA; Department of Molecular and Cellular Biology, The University of Arizona, Tucson, AZ 85721 USA

**Keywords:** RNA directed DNA Methylation, DNA methylation, CHH Methylation, Embryo development

## Abstract

**Background:** RNA directed DNA methylation (RdDM) initiates cytosine methylation in all contexts, and maintains asymmetric CHH methylation (where H is any base other than G). Mature plant embryos show one of the highest levels of CHH methylation, and it has been suggested that RdDM is responsible for this hypermethylation. Because loss of RdDM in *Brassica rapa* causes seed abortion, embryo methylation might play a role in seed development. RdDM is required in the maternal sporophyte, suggesting that small RNAs from the maternal sporophyte might translocate to the developing embryo, triggering DNA methylation that prevents seed abortion. This raises the question whether embryo hypermethylation is autonomously regulated by the embryo itself or influenced by the maternal sporophyte.

**Results:** Here, we demonstrate that *B. rapa* embryos are hypermethylated in both euchromatin and heterochromatin and that this process requires RdDM. Contrary to current models, *B. rapa* embryo hypermethylation is not correlated with demethylation of the endosperm. We also show that maternal somatic RdDM is not sufficient for global embryo hypermethylation, and we find no compelling evidence for maternal somatic influence over embryo methylation at any locus. Decoupling of maternal and zygotic RdDM leads to successful seed development despite loss of embryo CHH hypermethylation.

**Conclusions:** We conclude that embryo CHH hypermethylation is conserved, autonomously controlled, and not required for embryo development. Furthermore, maternal somatic RdDM, while required for seed development, does not directly influence embryo methylation patterns.

## Background

DNA methylation is an epigenetic modification that can modulate chromatin structure and gene expression [1]. Plants methylate cytosines in all sequence contexts (CG, CHG, and CHH, where H is any base other than G), and use specific methyltransferases to maintain each context after replication [2]. In addition, the RNA-directed DNA Methylation (RdDM) pathway is responsible for *de novo* methylation, a process that is most clearly observed at CHH positions [3]. RdDM functions primarily at the edges of euchromatin transposons, where constant re-establishment of methylation might be necessary [4,5].

RdDM can be divided into siRNA production and DNA methylation stages. During siRNA production, RNA Polymerase Pol IV produces single-stranded RNA transcripts which are copied into double-stranded RNA by RNA DEPENDENT RNA POLYMERASE 2 (RDR2) and cut into 24-nucleotide small interfering (si)RNAs by DICER LIKE 3 (DCL3) [6–8]. To mediate DNA methylation, these 24-nt siRNAs are loaded onto ARGONAUTE 4 (AGO4), which interacts with a non-coding scaffold transcript produced by RNA Polymerase V and recruits DOMAINS REARRANGED METHYLTRANSFERASE 2 (DRM2) to institute methylation marks on cytosine bases [9–11]. These two stages of RdDM frequently occur *in cis*, but can also function *in trans* due to siRNA-AGO4 loading in the cytoplasm [12]. SiRNAs can act *in trans* to trigger DNA methylation at allelic sites [13,14] or at homologous non-allelic sites [15], or might move between cells to act non-cell autonomously [16].

With the exception of Arabidopsis, which has only a small reduction in seed size, loss of RdDM in most species results in disruption of reproductive development, indicating that RdDM is necessary for successful sexual reproduction [17–21]. Mature embryos accumulate high levels of CHH methylation in Arabidopsis, soybean, and chickpea, suggesting that RdDM might enable reproduction through hypermethylation of the mature embryo [22–27]. In Arabidopsis, the developing endosperm is demethylated at sequences that show hypermethylation in the embryo, leading to the hypothesis that siRNAs produced in the endosperm might move to the embryo to direct methylation [22,28–30]. Movement of siRNAs between the maternal integuments and the filial tissues has also been proposed [31]. However, embryos produced through somatic embryogenesis also display hypermethylation, despite lack association with either endosperm or maternal integuments [27].

Here we show that *Brassica rapa* mature embryos are hypermethylated in the CHH context in both euchromatin and heterochromatin, and we demonstrate that this process requires RdDM. Although maternal RdDM is required for seed development, it is not sufficient for embryo hypermethylation, and methylation in the CHH context is not necessary for proper seed development. Furthermore, we find no evidence that hypermethylation of the embryo is driven by siRNAs produced in adjacent tissues, suggesting that embryo CHH hypermethylation is entirely autonomous.

## Results

### *Brassica rapa* embryos are hypermethylated in CHH Context

To analyze global methylation levels in mature embryos, we performed whole-genome bisulfite sequencing on embryos dissected from dry seeds, and compared the resulting data with other reproductive tissues (ovule, endosperm, and early embryo) and a non-reproductive control (leaf). Bisulfite conversion was greater than 99% in all samples with read depth coverage > 9 (Supplemental Table 1). We calculated methylation in 300-bp non-overlapping windows for all three sequence contexts (Figure 1A). CG methylation was largely unchanged across the tissues, with the exception of endosperm, which was demethylated in CG and CHG contexts, consistent with observations in Arabidopsis and rice [32–34]. As in Arabidopsis, soybean, and chickpea [22–27] we also observed elevated global CHH methylation in *B. rapa* mature embryos, with moderately increased CHH methylation in torpedo-stage embryos (Figure 1A). The increased CHH methylation in torpedo-stage embryos was correlated with CHH hypermethylation in mature embryos (Figure 1B, correlation coefficient = 0.6), indicating that hypermethylation is a gradual process during embryogenesis.

**Figure 1.**
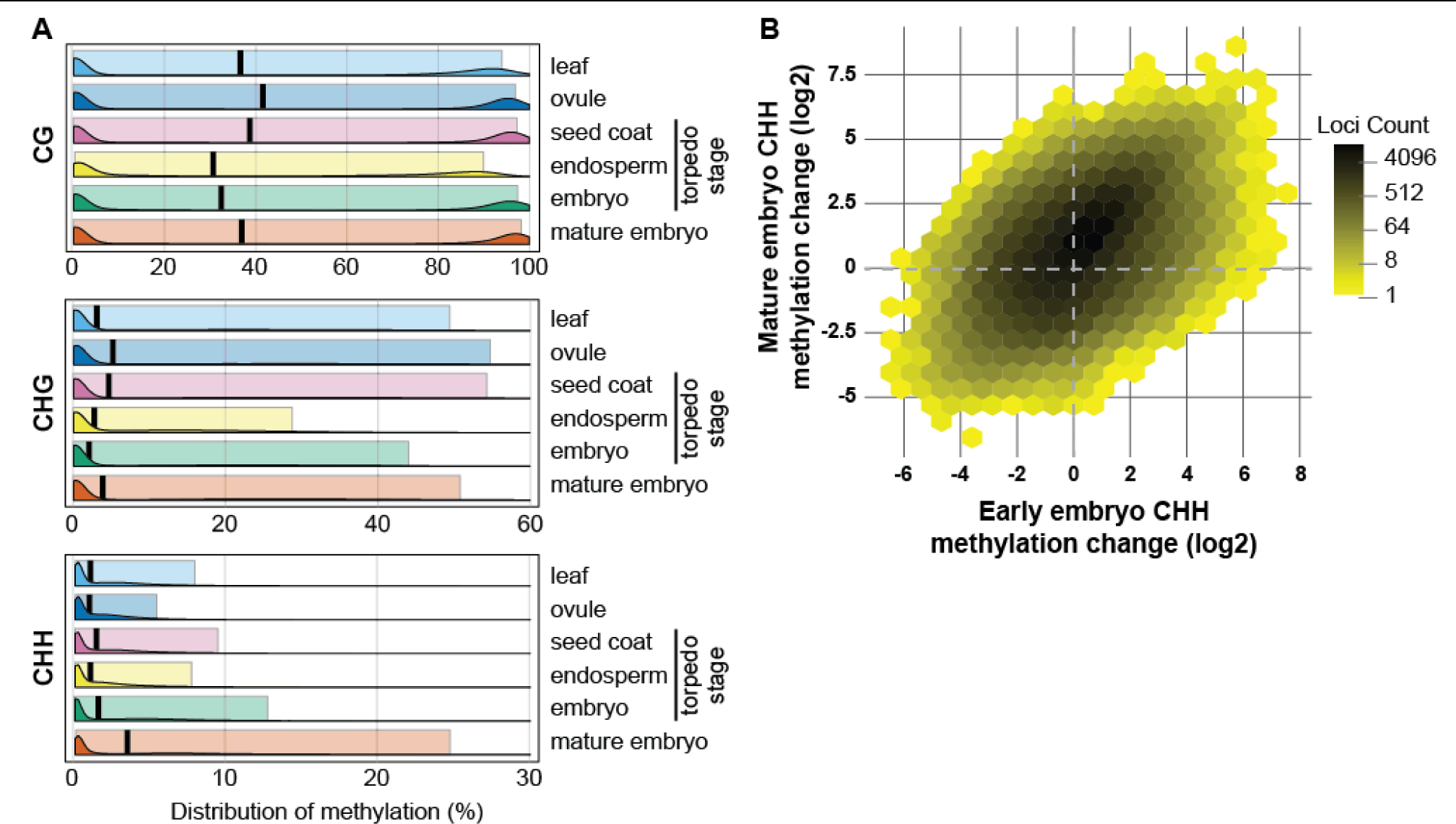
*B. rapa* embryos are hypermethylated at CHH sites. (A) Distribution of methylation levels in 300-nt windows across the *B. rapa* genome. Ridge plots display density of average methylation in each context, while background box plots enclose the 10^th^ to 90^th^ percentiles of the data. The black bar marks the median for each tissue/context combination. Only windows with read depth ≥ 5 over all cytosines were included (approximately 1 million windows per tissue/context combination). (B) Increased CHH methylation (log_2_ fold-change relative to leaves) is correlated in torpedo and mature embryos. 752,405 300-nt windows with read depth of at least 5 in both tissues are plotted.

To assess the types of chromatin responsible for embryo hypermethylation, we analyzed methylation levels in 25-kb windows across each chromosome (Figure 2A and Supplemental Figure 1). Pericentromeric heterochromatin, which has denser accumulation of transposons, was strongly methylated in CG and CHG contexts in all tissues. The sole exception was endosperm, which as expected showed a small reduction in CG methylation and stronger loss of CHG methylation. In comparison to these heterochromatic marks, CHH methylation was distributed more equally across the length of the chromosome. Increased CHH methylation in mature embryos relative to other tissues was readily apparent. In Arabidopsis, embryo hypermethylation occurred primarily in pericentromeric heterochromatin [22,23], while in soybean somatic embryos, CHH hypermethylation was seen across the genome [27]. We observed a similar degree of CHH hypermethylation in both the CG-dense pericentromeric regions and the chromosome arms (Figure 2B), indicating that both heterochromatin and euchromatin are targets of CHH methylation during *B. rapa* embryo development.

**Figure 2.**
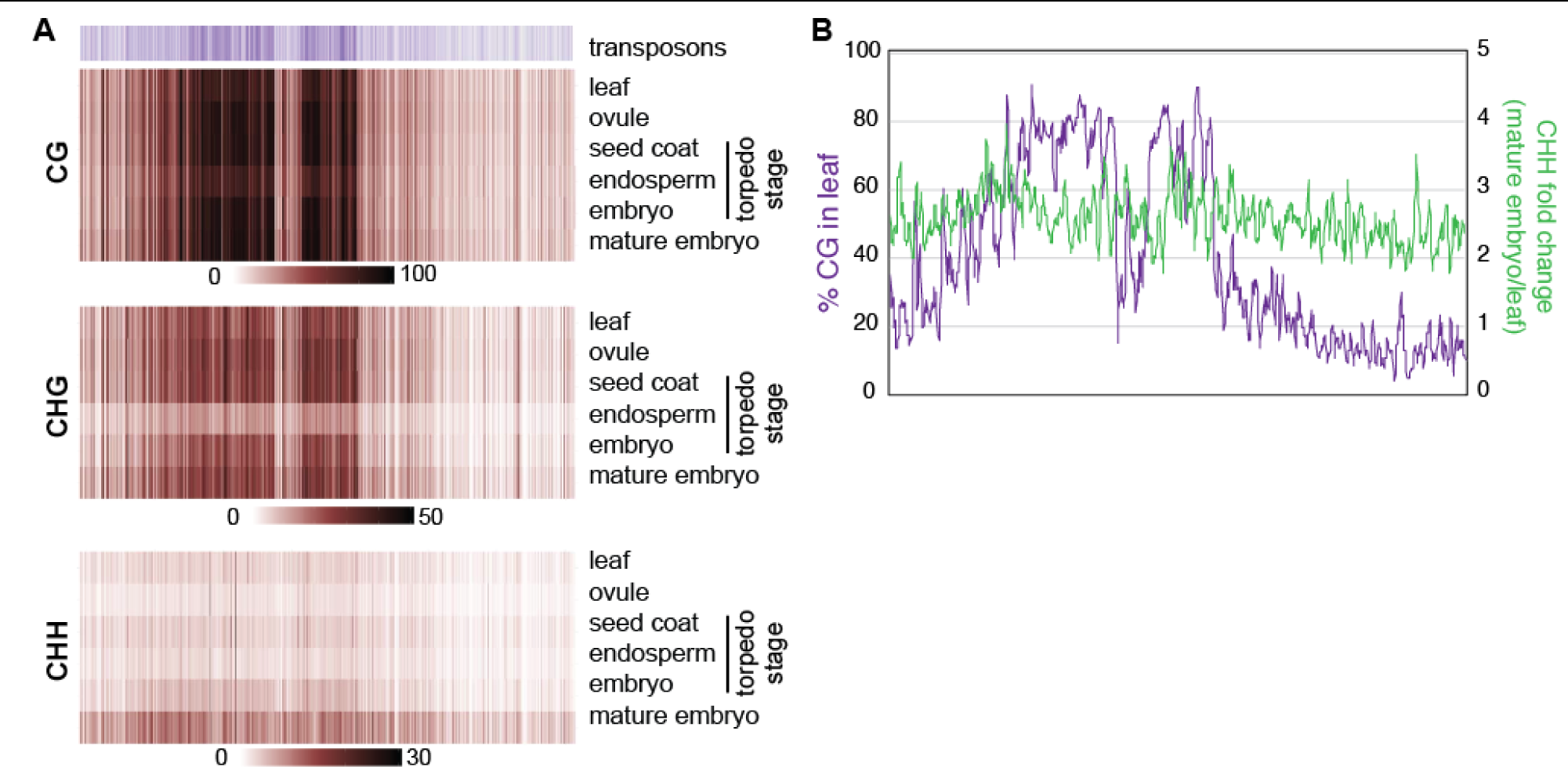
Embryo CHH hypermethylation is not restricted to the pericentromere. (A) Heatmaps of transposon density or methylation level in 25-kb windows across chromosome 10. Each methylation context has its own scale bar to visualize changes across tissues. Other chromosomes are presented in Supplemental Figure 1. (B) CHH hypermethylation in mature embryos (green line) is not correlated with the amount of CG methylation in leaves (purple line). Five-window rolling average of 25-kb windows across chromosome 10 are plotted.

Finally, we assessed embryo methylation relative to leaf for each cytosine context within each 300-nt window. Most windows were unchanged with respect to CG and CHG methylation, while CHH methylation showed a pronounced shift toward hypermethylation (Figure 3A). These changes were highly significant (Figure 3B), but we selected only windows with the strongest changes in methylation for further analysis. We defined differentially methylated windows (DMWs) as those with at least 5-fold (log_2_ = 2.32) increase or decrease in methylation in mature embryo compared to leaf, and an FDR adjusted p-value less than 0.005 (Figure 3A, 3B). Hypermethylated windows were more abundant than hypomethylated windows for each sequence context, with CHH hyper-DMWs vastly outnumbering other DMWs in other sequence contexts (Figure 3C).

**Figure 3.**
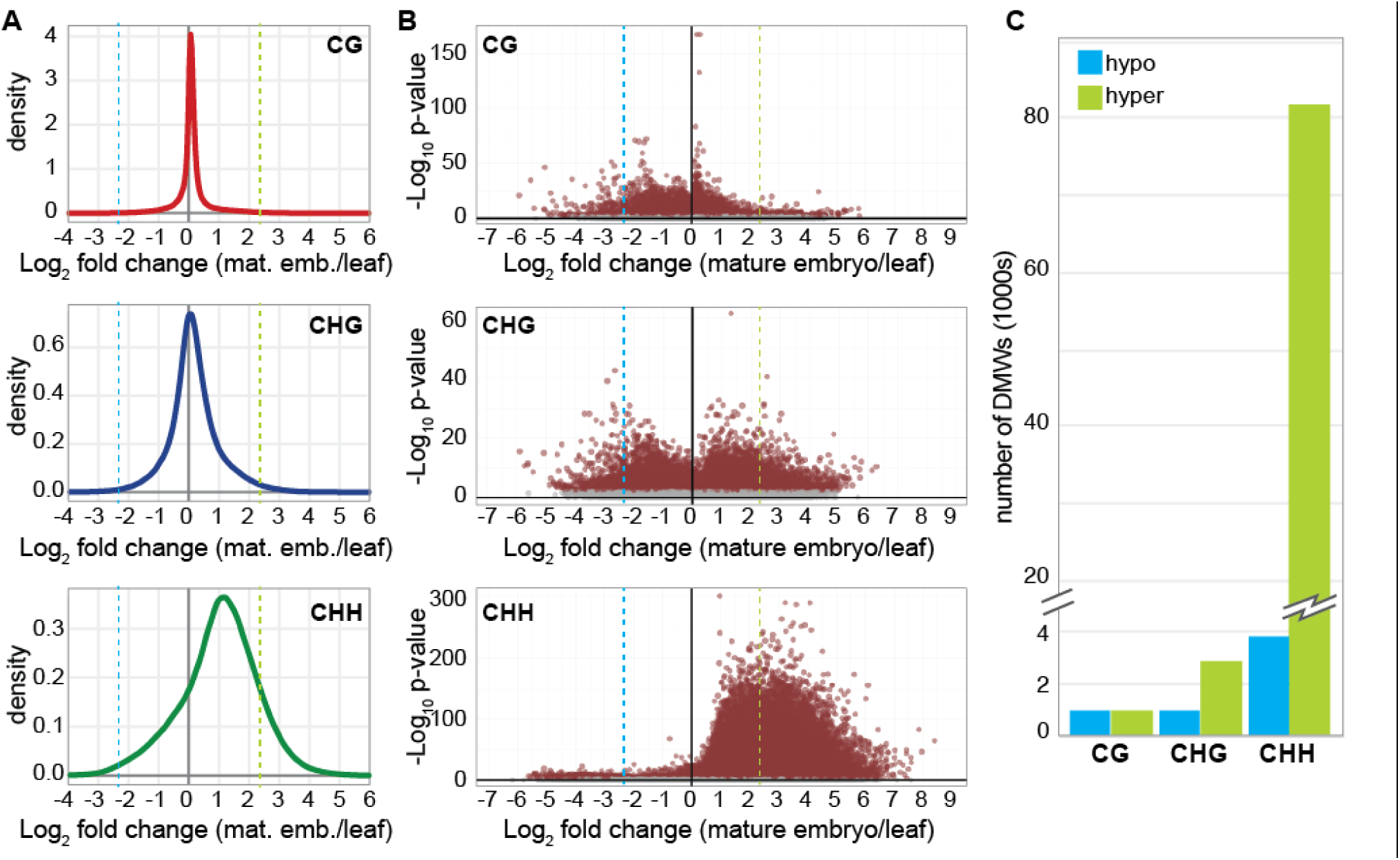
Identification of embryo Differentially Methylated Windows. Density distributions (A) and volcano plots (B) of methylation fold change in mature embryos versus leaf for three cytosine contexts. 300-nt windows with read depth of at least 5 are plotted (650,665 CG, 686,741 CHG, and 869,526 CHH windows). The dashed blue line marks 5-fold hypomethylation and the dashed green line marks 5-fold hypermethylation. Windows above this threshold with an FDR-adjusted p-value <0.005 were collected for subsequent DMW analysis. (C) Number of differentially methylated windows (DMWs) passing the above thresholds in each methylation context.

Together, our observations demonstrate that mature *B. rapa* embryos are extensively hypermethylated at CHH sites across the genome, and this hypermethylation is the primary difference between leaf and embryo methylation patterns. This hypermethylation is widespread, not limited to pericentromeric heterochromatin, and progressive throughout embryogenesis.

### Embryo CHH hypermethylation is dependent on RdDM

RdDM is the major pathway for *de novo* methylation in all sequence contexts, and its activity is frequently observed through accumulation of CHH methylation. However, most of the CHH methylation in the genome is instead placed by CHROMOMETHYLTRANSFERASE 2 (CMT2) [18,35]. Kawakatsu and colleagues [23] demonstrated that both RdDM and CMT2 contribute to CHH methylation in the embryo, but their analysis did not determine which process was responsible for the hypermethylation relative to non-embryonic tissues. Small RNA accumulation at hypermethylated regions is correlated with embryo hypermethylation [22,24,26] but it is not clear whether siRNA accumulation is required for increased methylation, or whether these two processes occur independently.

Our analysis also implicates RdDM in embryo CHH hypermethylation. Firstly, compared to all genomic windows with sufficient WGBS read depth, embryo CHH hyper-DMWs are significantly enriched for Class II DNA elements and Class III Helitrons (Figure 4A). CHH hyper-DMWs are also significantly depleted at genes and at long terminal repeat (LTR) retroelements, following the characteristic pattern of RdDM loci in *B. rapa* [19]. Most importantly, CHH hyper-DMWs are enriched for loci previously shown to produce 24-nt siRNAs (Figure 4A).

**Figure 4.**
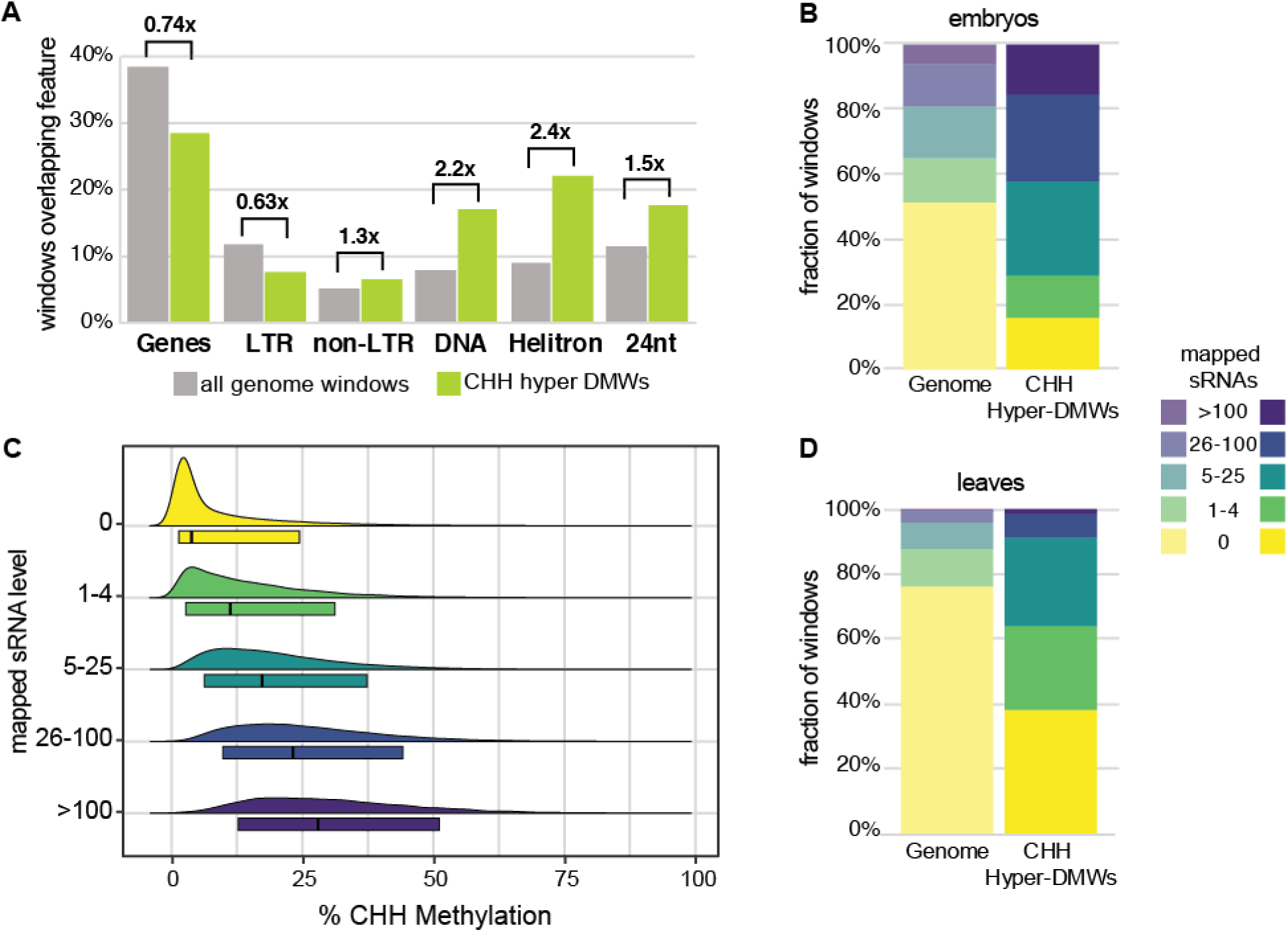
small RNA production, but not abundance, is correlated with embryo hypermethylation. (A) Enrichment or depletion of genomic features at CHH hyper-DMWs. The percentage of CHH hyper-DMWs overlapping annotated genomic features is plotted compared to the percentage of overlap for all genomic windows with comparable read depth. All differences are significant at p<2.2e-16. (B, D) CHH hyper-DMWs and all genomic windows were binned based on the number of sRNAs mapping to them in torpedo embryos or leaves, and the fraction of windows in each bin is shown. The embryo sRNA library has 39.1 million mapped reads, while the leaf library has 10.8 million reads. (C) Absolute CHH methylation in mature embryos is plotted as a function of the number of mapped sRNAs in torpedo embryos at all CHH hyper-DMWs. Box plots circumscribe the 10^th^-90^th^ percentiles and the black bar marks the median.

To further investigate the association between CHH hyper-DMWs and siRNAs, we analyzed siRNA accumulation at CHH hyper-DMWs in torpedo-stage embryos (Supplemental Table 2). CHH hyper-DMWs have significantly higher accumulation of siRNAs than the genome as a whole (Figure 4B) and windows with greater siRNA accumulation in torpedo embryos have higher CHH methylation in mature embryos (Figure 4C). However, many CHH hyper-DMWs lack substantial siRNA accumulation, a pattern also detected in chickpea [26]. We also observed a similar enrichment of siRNAs at CHH hyper-DMWs in leaves despite the 5-fold or greater difference in methylation between these tissues (Figure 4D).

To directly test whether RdDM is required for embryo hypermethylation in *B. rapa*, we assayed differences in methylation levels at CHH hyper-DMWs between wild type and an RdDM-deficient mutant, *braA*.*rdr2-2* (*rdr2* hereafter) (Grover *et al*., 2018). Mature *rdr2* embryos have a clear reduction in CHH and CHG methylation compared to wild-type embryos (Figure 5A, B), with CHH methylation levels similar to wild-type leaves. In contrast, CG methylation at these CHH hyper-DMWs is *rdr2*-independent. This pattern clearly demonstrates that embryo hypermethylation in the CHH context is due to RdDM, and that other methylation pathways have only a minor contribution to embryo CHH hypermethylation.

**Figure 5.**
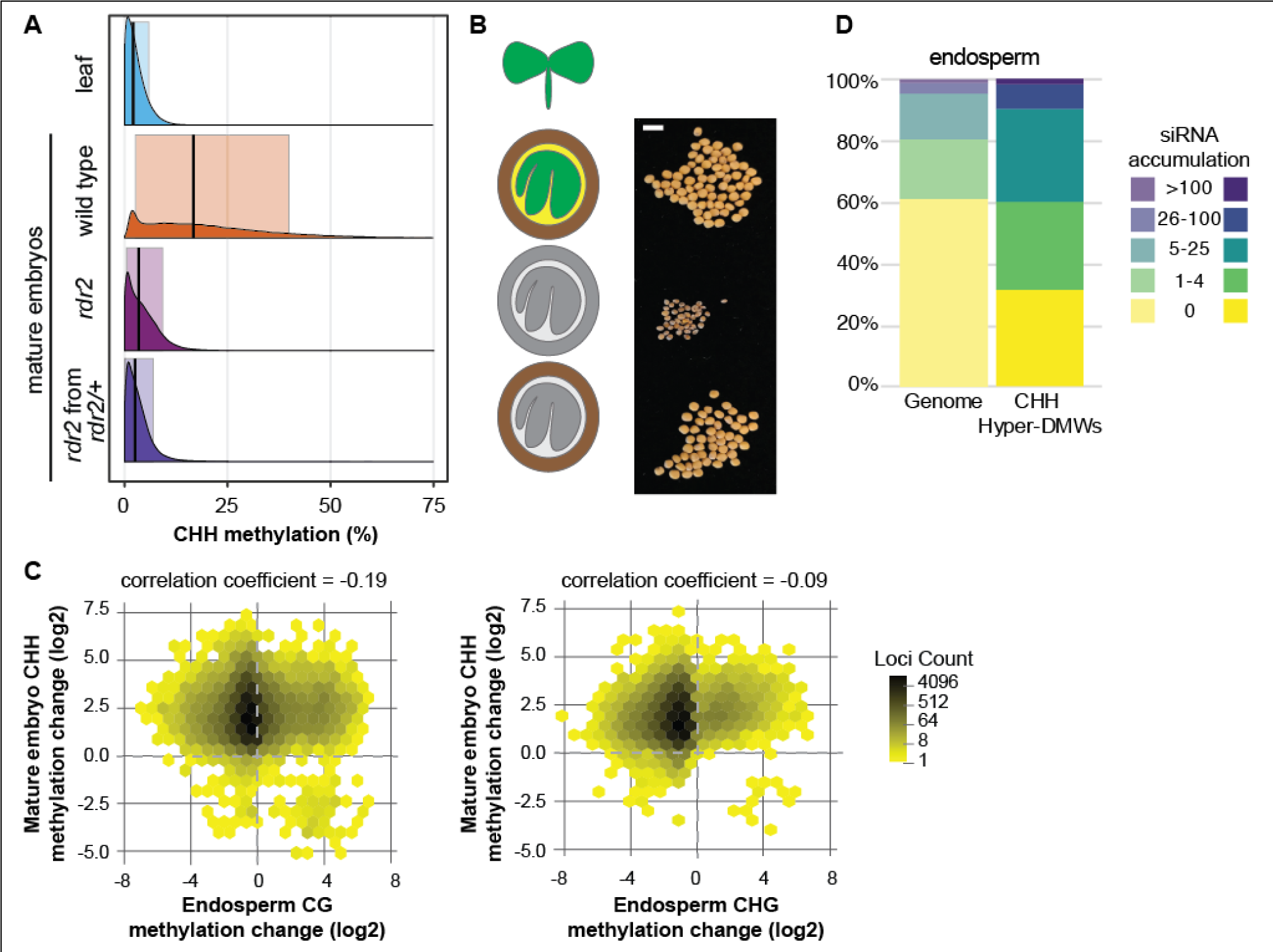
Embryo hypermethylation is determined by filial genotype. (A) Distribution of CHH methylation at CHH hyper-DMWs in leaves and mature embryos. *rdr2* embryos were derived either from *rdr2* homozygous mothers (maroon) or from *rdr2/+* heterozygous mothers (purple). (B) Cartoon and images of representative seeds measured in A. Colored tissues have functional RdDM; grey tissues are deficient in RDR2. Scale bar is 5 mm. (C) Hex plots of mature embryo CHH methylation change by torpedo-stage endosperm CG (left) or CHG (right) methylation. (D) siRNA accumulation in endosperm at CHH hyper-DMWs. The endosperm siRNA library has 19.6 million mapped reads.

### No evidence for endosperm-directed hypermethylation of the embryo

It has been suggested that demethylation of the endosperm allows production of siRNAs that target DNA methylation in the embryo [22,28–30]. To determine whether there is an association between endosperm demethylation and embryo hypermethylation, we compared changes in methylation levels in these tissues. Compared to leaf samples, endosperm is demethylated for both CG and CHG, while embryos are hypermethylated for CHH (Figure 1A). However, there is no correlation between CG or CHG demethylation in the endosperm and CHH hypermethylation in the embryo, whether we assessed all genomic windows (Figure 5C) or only the CHH hyper-DMWs (Supplemental Figure 2). We measured correlations between embryo and endosperm methylation in multiple ways, both globally and at CHH hyper-DMWs (Supplemental Table 3). Absolute CHH methylation in the embryo is *positively* correlated with all methylation contexts in the endosperm, indicating that the embryo hypermethylated loci tend to have higher absolute methylation in endosperm. When fold change in methylation relative to leaf samples are measured, only CHH methylation is correlated between these tissues, indicating that regardless of their absolute methylation level, the filial tissues are coordinately increasing CHH methylation. Only when we use difference in absolute methylation as a metric do we find a correlation between endosperm CG/CHG demethylation and embryo CHH hypermethylation. However, this correlation is not strong and become weaker when only the CHH hyper-DMWs are assessed. Together, these results suggest that any correlation between endosperm demethylation and embryo CHH hypermethylation is not robust to analysis method and is likely artifactual.

Because the presumed mechanism whereby endosperm demethylation triggers embryo hypermethylation is transport of siRNAs, we also assessed siRNA production at CHH hyper-DMWs in developing endosperm (Figure 5D). CHH hyper-DMWs produce more siRNAs than the genome as a whole, but at a level that is comparable to developing embryos or leaves (Figure 4B, 4D), suggesting that siRNA production occurs at these windows consistently across tissues and is not a response to endosperm demethylation. On the whole, we find no evidence that demethylation of the endosperm triggers siRNA production to cause hypermethylation of the mature embryo.

### Maternal sporophytic RdDM is not sufficient for embryo hypermethylation

RdDM mutants in *Capsella rubella* and *Brassica rapa* show a high rate of seed abortion that is dependent on the maternal somatic genotype rather than the filial genotype [19,36]. The few seeds that are produced from *rdr2* plants are smaller and irregular in size and shape (Figure 5E). Because a functional RdDM pathway in the maternal sporophyte is required for seed development, we hypothesized that maternal sporophytic RdDM might drive hypermethylation of the developing embryo. To test this hypothesis, we pollinated heterozygous (*rdr2/RDR2*) pistils with homozygous (*rdr2/rdr2*) pollen and identified *rdr2* embryos that developed in the presence of functional maternal sporophytic RdDM. Compared to methylation levels of *rdr2* mutant embryos from homozygous mutant mothers (*rdr2/rdr2*), we did not observe restoration of embryo CHH hypermethylation (Figure 5A, B).

To further probe the possibility that maternal sporophytically-derived siRNAs might trigger hypermethylation of the embryo, we assessed DNA methylation levels at previously-defined “siren” loci [31]. These loci produced over 90% of the 24-nt siRNAs in maternal integuments and are also the most highly-expressed siRNA loci in endosperm. We found no evidence that siRNAs produced from siren loci in the integument were able to direct DNA methylation in *rdr2* embryos (Supplemental Figure 3). These results indicate that although siRNA production in the maternal sporophyte is necessary for seed development, it is not sufficient for embryo hypermethylation.

Together these observations provide no evidence that embryo hypermethylation is directed by siRNAs from another tissue. Combined with the observation that embryos formed through somatic embryogenesis also have elevated DNA methylation despite their lack of interaction with endosperm or integuments, we conclude that embryo hypermethylation is autonomously directed.

## Discussion

Seeds form the majority of the world’s food supply, making the development of the seed and interactions between its the multiple tissues critically important areas for research. Double fertilization gives rise to the diploid embryo and the triploid endosperm, which are surrounded by the seed coat, a maternal somatic tissue. Communication between maternal and filial tissues, as well as between the embryo and endosperm, is essential to coordinate the development of a seed [37,38]. Small RNAs have been proposed to move between seed tissues and to establish robust methylation of transposons at this transition between generations [22,28–31].

Here, we provide direct evidence that 24-nt siRNAs are responsible for hypermethylation of mature embryos by demonstrating that *rdr2* embryos lose hypermethylation. However, our evidence suggests that these siRNAs are derived autonomously in the embryo and are not transported from other tissues. Maternal sporophytic *RDR2* (and hence, siRNA production) is not sufficient for embryo hypermethylation (Figure 5A, B), clearly indicating that the siRNAs responsible for hypermethylation are produced in the filial tissues. Because the embryo and the endosperm have the same genotype (differing only in maternal ploidy), we cannot separate them genetically. However, we find that endosperm does not produce more siRNAs than embryos from CHH hyper-DMWs (Figure 5D), nor is there correlation between endosperm CG/CHG demethylation and embryo CHH hypermethylation (Figure 5C). Furthermore, somatic soybean embryos produced in tissue culture also display embryo hypermethylation [27]. The most parsimonious explanation for these observations is that embryo CHH hypermethylation is autonomously directed by siRNAs synthesized in the embryo.

Although maternal sporophytic siRNA production is not sufficient for embryo hypermethylation, it remains possible that siRNAs from the maternal integument might trigger the expression of 24-nt siRNAs in the embryo and initiate autonomous methylation in the embryo. This process would be analogous to the production of piRNAs in *Drosophila melanogaster*, whereby maternally-derived small RNAs initiate subsequent filial siRNA production and transposon silencing [39]. It also remains possible that triggering siRNAs are brought to the zygote during fertilization by the sperm nucleus, although this model remains to be tested [40,41].

Despite a lack of embryo hypermethylation, *rdr2* homozygous seeds from heterozygous mothers phenocopy wild-type seeds (Figure 5E), indicating that embryo hypermethylation is not necessary for seed development in *B. rapa*. Similarly, Arabidopsis does not require DRM2 methyltransferase for embryo development [25], however Arabidopsis does not require RdDM generally, while other species in Brassicaceae have reproductive defects in the absence of RdDM [19]. Decoupling of embryo methylation and seed development in *B. rapa* supports the hypothesis that embryo hypermethylation is important for seed dormancy or longevity but not for seed development [22–24]. We assessed segregating seed populations and observed no difference in germination timing or frequency for unmethylated *rdr2* embryos relative to their methylated siblings (data not shown), suggesting that other hypotheses should also be considered.

In Arabidopsis, embryo hypermethylation is preferentially targeted to transposons in the pericentromeric heterochromatin [22,23], while hypermethylation also occurs at euchromatic transposons in soybean [25,27]. In *B. rapa*, we detect hypermethylation in both heterochromatin and euchromatin (Figure 2B), suggesting that euchromatic embryo hypermethylation might be common among plants. Recent work demonstrates that Arabidopsis heterochromatin is decondensed and produces abundant 24-nt siRNAs during embryogenesis [42], providing an opportunity for the RdDM machinery to access this chromatin for hypermethylation.

We were surprised by the low level of siRNAs at CHH hyper-DMWs in torpedo-stage embryos. Correlation between CHH levels in torpedo-stage and mature embryos (Figure 1B) indicates that hypermethylation occurs throughout embryogenesis rather than during embryo maturation, and therefore robust siRNA accumulation would be predicted. The 81,556 CHH hyper-DMWs account for ∼10% of windows with sufficient read depth, and they accumulate 16.9% of the mapped siRNAs (20.1% of the mapped 24-nt siRNAs). While this is a substantial enrichment compared to the genome as a whole, these windows account for 13.8% of the mapped siRNAs (15.9% of the mapped 24-nt siRNAs) in leaves. This discrepancy suggests that while siRNA production is required for embryo hypermethylation, developmental-specific factors are required for robust methylation.

## Conclusions

*Brassica rapa* embryos are hypermethylated at both euchromatic and heterochromatic CHH positions. This hypermethylation requires RdDM, and there is no evidence that siRNAs from the endosperm or maternal somatic tissue direct embryo methylation. Successful development of seeds lacking embryo hypermethylation indicate that this methylation is not necessary for embryogenesis, even in species that require RdDM for seed development.

## Methods

### Plant materials and growth conditions

*Brassica rapa ssp trilocularis* variety R-o-18 were grown in a greenhouse at 70°/60°F (day/night) under at least 16 hours of illumination. Plants were fully dried before seed collection. Dry seeds were soaked in water for no more than 60 minutes before manual dissection to remove mature embryos. Three wild-type or five *rdr2* mutant embryos were pooled before DNA extraction with the GeneJET Plant Genomic DNA Purification Kit (Thermo Fisher Scientific, K0791). Embryos from *rdr2/RDR2* heterozygous mothers were individually collected, prepped, and genotyped prior to DNA pooling. Torpedo-stage endosperm and embryo samples were dissected from pistils that were manually pollinated with *B. rapa* genotype R500. Whole-genome bisulfite sequencing libraries were prepared as previously described [43]. Lambda Phage DNA (Promega D1521) was included as a bisulfite conversion control. Libraries were pooled and sequenced in a single lane of paired-end 76-nt on an Illumina NextSeq500 at the University of Arizona Genetics Core.

### Methylation Analysis

Whole Genome Bisulfite Sequencing data from ovule and leaf were obtained from NCBI (BioProject PRJNA588293). For other tissues, sequencing reads were quality controlled with FastQC [44,45] and trimmed using Trim Galore (options --trim-n and --quality 20) [46]. Trimmed reads were aligned to *Brassica rapa* R-o-18 genome (v2.2, a kind gift from G.J. King and the *B. rapa* sequencing consortium) with bwameth [47]. To mark PCR duplicates and determine properly paired alignment rate, Picard Tools [48] and Samtools [49] were respectively used, with options -q 10, -c, -F 3840, -f 66 for Samtools. We used Mosdepth [50] with option -x and -Q 10 and a custom Python script developed previously in the lab (bed_coverage_to_x_coverage.py, https://github.com/The-Mosher-Lab/grover_et_al_sirens_2020) to determine genomic coverage. Statistics for all libraries are found in Supplemental Table 1.

Percentage methylation per cytosine was extracted with MethylDackel [51] in two successive steps. The first step was to identify inclusion bounds based on methylation bias per read position using MethylDackel mbias, followed by MethylDackel extract. Since the default for MethylDackel is the CG context, we also used --CHG and --CHH options. We determined bisulfite conversion rates by alignment to the bacteriophage lambda (NCBI Genbank accession J02459.1) and *Brassica rapa* var. *pekinensis* chloroplast (NCBI Genbank accession NC 015139.1) genomes with a custom Python script developed previously in the lab (bedgraph_bisulfite_conv_calc.py, https://github.com/The-Mosher-Lab/grover_et_al_sirens_2020). Conversion frequencies were all above 99.4% (Supplemental Table 1). Replicates were checked for consistency by principal component analysis before pooling to increase read depth (Supplemental Figure 4).

Pairwise methylation difference analysis between tissues were done using the methylKit package and a custom script with window size of 300, and with a q value of 1000 with percentage methylation difference of zero to obtain all data across all windows in the genome and filter it with R. Methylation was calculated for each sample on 300-bp non-overlapping windows, which was made with the help of BEDTools makewindows [52] feature on the *Brassica rapa* R-o-18 genome.

We calculated enrichment of genomic features like small RNA loci, TEs, and genes within the CHH hyper-DMWs using BEDTools intersect [52,53]. Transposable elements were annotated as in [31]. We considered features to be overlapping if there was at least 1 nucleotide shared. Genomic features were annotated onto the 300-bp non-overlapping windows using BEDTools makewindows [52] and the number of overlaps and non-overlaps between the hyper-DMWs and the genomic features were recorded. A Fisher’s exact test was performed in R to determine if the number of overlaps indicated significant enrichment or depletion.

Methylation over pre-defined siren loci [31] was determined with BEDTools intersect [52] and a custom Python script [(bedgraph_methylation_by_bed.py) developed previously [31] in the lab.

### Small RNA analysis

Small RNA sequencing datasets were obtained from NCBI (BioProject PRJNA588293, Supplemental Table 2). Small RNA processing (quality checking, noncoding RNA filtering, removal of reads mapping to chloroplast and mitochondrial genomes) was carried out with a publicly-available small RNA data processing pipeline [54]. Only 19 to 26-nt reads were retained for further analysis. Replicates were pooled for better read alignment and depth. The genome was divided into 300-bp non-overlapping windows using BEDTools makewindows [52] and ShortStack [55,56] was used to get read counts on each window (options --mismatches 0, --mmap u, --mincov 0.5 rpm, --pad 75 and --foldsize 1000). The sum of all 19-26nt small RNA reads from genomic windows or CHH hyper-DMWs were low and susceptible to count-based bias when normalized against total library size. Therefore, windows were binned into 5 sRNA expression levels and compared only within the same tissue.

## Declarations

### Availability of data and materials

The datasets supporting the conclusions of this article are available in the National Center for Biotechnology Information Sequence Read Archive under BioProjects PRJNA588293 and PRJNA657007.

### Authors’ contributions

RAM conceptualized the project; TC, TK, and JWG performed experiments; TC and RAM analyzed and visualized the data and wrote the paper.

## Acknowledgements

The authors are grateful to The *Brassica rapa* R-o-18 Genome Sequencing Consortium for access to the *B. rapa* v2.3 genome and annotations and to Dr. Diane Burgess for help with transposable element annotation. Illumina sequencing was performed by The University of Arizona Genetics Core.

---

**Supplemental Figure 1.**
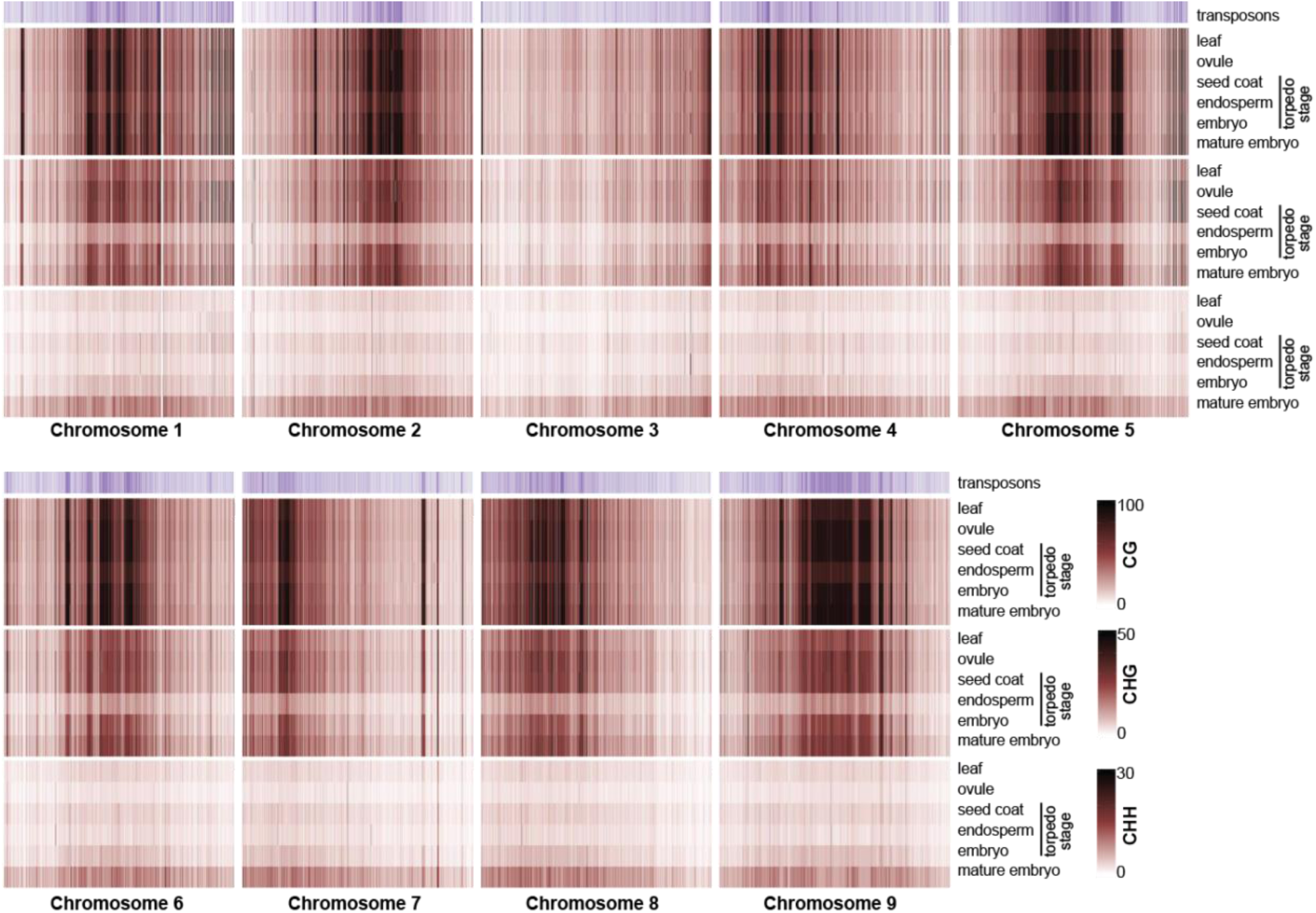
Heatmaps of methylation on *B. rapa* chromosomes 1-9. Heat maps of methylation in 25-kb windows across *B. rapa* chromosomes, as in Figure 2A.

**Supplemental Figure 2.**
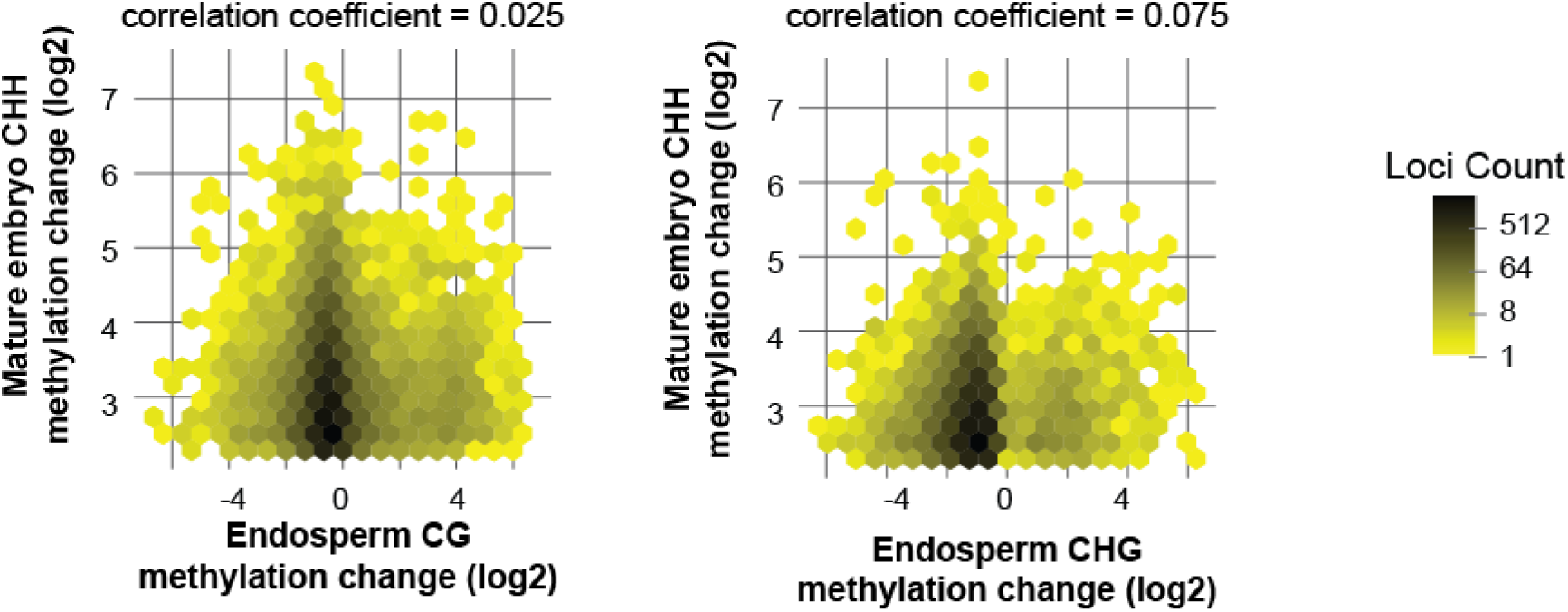
No correlation between endosperm demethylation and embryo hypermethylation at CHH hyper-DMWs. Hex plots of mature embryo CHH methylation change by torpedo-stage endosperm CG (left) or CHG (right) methylation change. Only CHH hyper-DMWs are plotted (whole genome plotted in Figure 5C).

**Supplemental Figure 3.**
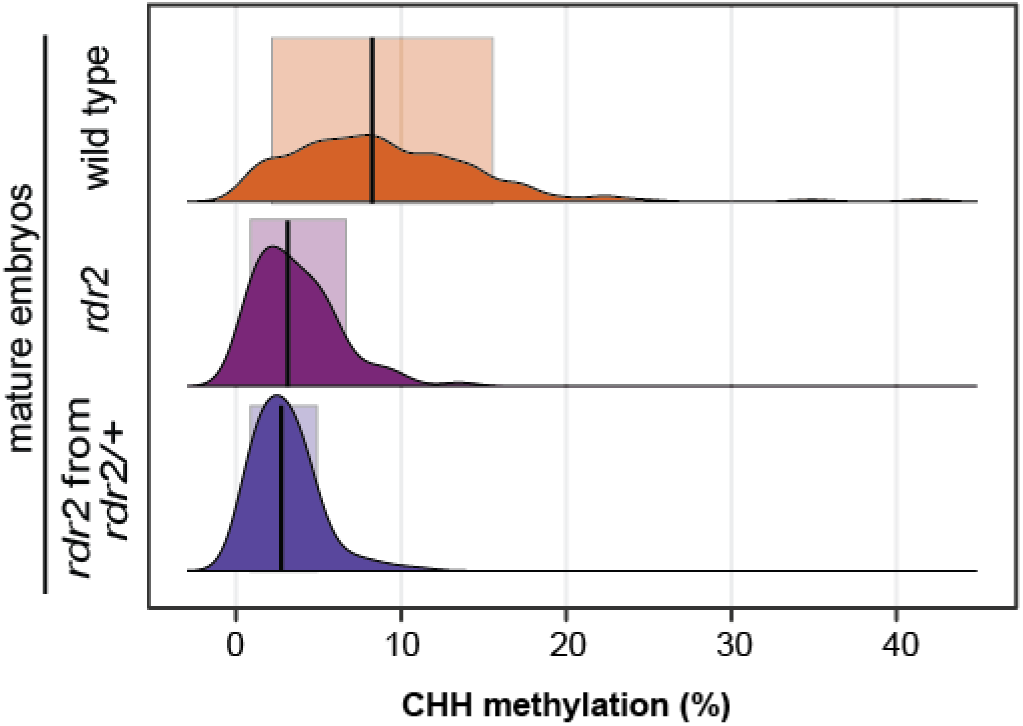
Mature embryo methylation at siren loci. Distribution of CHH methylation at siren loci in mature embryos. *rdr2* embryos were derived either from *rdr2* homozygous mothers (maroon) or from *rdr2/+* heterozygous mothers (purple).

**Supplemental Figure 4.**
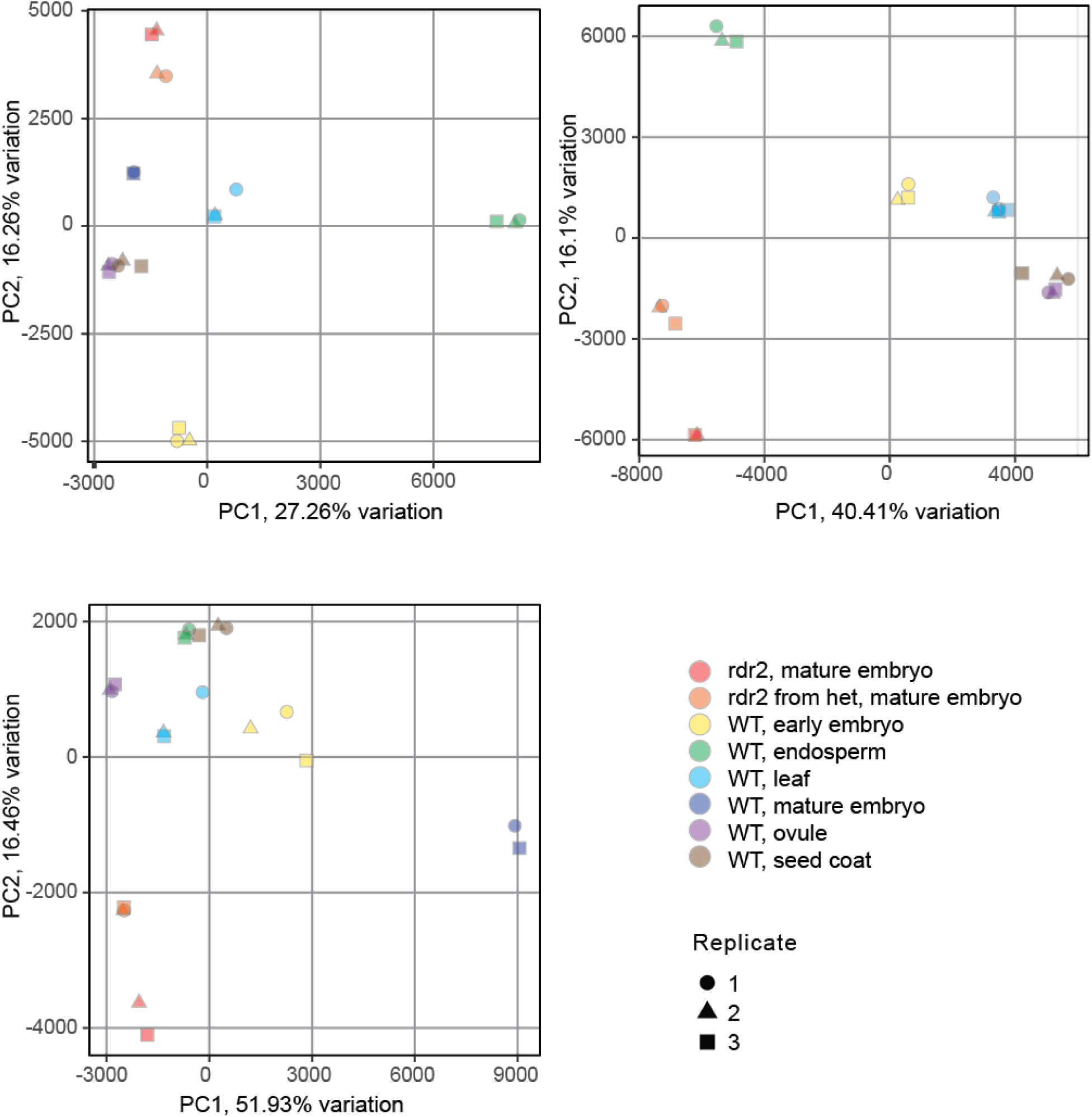
PCA analysis of WGBS replicates.

**Supplemental Table 1.** WGBS sequence datasets.

| Genotype | Tissue | Replicate | NCBI identifier | De-multiplexed Reads | Deduplicated Alignments | Mapping Rate % | Coverage X | Conversion Rate (lambda DNA) | Conversion Rate (chloroplast DNA) |
| --- | --- | --- | --- | --- | --- | --- | --- | --- | --- |
| WT R-o-18 | leaf | 1 | SRX7113698 | 33,843,130 | 18,626,017 | 55.43% | 8.24 | 99.66 | 99.60 |
| WT R-o-18 | leaf | 2 | SRX7113699 | 34,414,305 | 16,928,242 | 49.56% | 7.46 | 99.68 | 99.66 |
| WT R-o-18 | leaf | 3 | SRX7113700 | 39,216,330 | 19,653,280 | 50.69% | 8.73 | 99.63 | 99.61 |
| WT R-o-18 | ovule | 1 | SRX7113704 | 35,677,253 | 20,580,397 | 58.35% | 9.08 | 99.64 | 99.52 |
| WT R-o-18 | ovule | 2 | SRX7113705 | 38,415,932 | 22,250,022 | 58.58% | 9.81 | 99.65 | 99.51 |
| WT R-o-18 | ovule | 3 | SRX7113706 | 33,159,726 | 19,047,779 | 58.06% | 8.38 | 99.63 | 99.47 |
| WT R-o-18 x R500 | endosperm | 1 | SRX8941582 | 37,703,830 | 20,112,987 | 54.30% | 9.09 | 99.71 | 99.55 |
| WT R-o-18 x R500 | endosperm | 2 | SRX8941583 | 40,064,286 | 20,957,711 | 53.18% | 9.61 | 99.69 | 99.49 |
| WT R-o-18 x R500 | endosperm | 3 | SRX8941584 | 42,843,140 | 22,831,013 | 53.88% | 10.40 | 99.70 | 99.50 |
| WT R-o-18 x R500 | seed coat | 1 | SRX8941585 | 37,809,229 | 20,014,869 | 53.89% | 9.10 | 99.71 | 99.36 |
| WT R-o-18 x R500 | seed coat | 2 | SRX8941586 | 38,519,379 | 19,894,934 | 50.06% | 8.94 | 99.76 | 99.43 |
| WT R-o-18 x R500 | seed coat | 3 | SRX8941587 | 40,731,277 | 21,863,906 | 54.61% | 9.95 | 99.69 | 99.36 |
| WT R-o-18 x R500 | torpedo embryo | 1 | SRX8941594 | 37,095,900 | 18,335,536 | 49.81% | 5.81 | 99.72 | 99.65 |
| WT R-o-18 x R500 | torpedo embryo | 2 | SRX8941595 | 37,168,809 | 18,136,734 | 50.34% | 8.35 | 99.71 | 99.66 |
| WT R-o-18 x R500 | torpedo embryo | 3 | SRX8941581 | 39,057,700 | 19,063,283 | 49.35% | 8.94 | 99.67 | 99.63 |
| rdr2 R-o-18 | mature embryo | 1 | SRX8941580 | 42,990,881 | 22,738,273 | 52.89% | 10.02 | 99.57 | 99.56 |
| rdr2 R-o-18 | mature embryo | 2 | SRX8941588 | 44,373,420 | 24,006,617 | 54.10% | 10.51 | 99.52 | 99.48 |
| rdr2 from het mother R-o-18 | mature embryo | 1 | SRX8941589 | 48,591,584 | 24,052,250 | 49.50% | 10.83 | 99.62 | 99.57 |
| rdr2 from het mother R-o-18 | mature embryo | 2 | SRX8941590 | 45,727,329 | 22,713,974 | 49.67% | 10.11 | 99.56 | 99.54 |
| rdr2 from het mother R-o-18 | mature embryo | 3 | SRX8941591 | 45,076,722 | 21,617,299 | 47.96% | 9.77 | 99.59 | 99.56 |
| WT R-o-18 | mature embryo | 1 | SRX8941592 | 48,525,785 | 24,826,190 | 51.16% | 10.95 | 99.43 | 99.52 |
| WT R-o-18 | mature embryo | 2 | SRX8941593 | 40,351,733 | 20,545,946 | 50.92% | 9.04 | 99.50 | 99.57 |

**Supplemental Table 2.** sRNA sequence datasets.

| Genotype | Tissue | Replicate | NCBI identifier | De-multiplexed Reads | Trimming & q > 30 filtering | % Remaining | rfam *B. rapa* & *A. thaliana* Filtered | % Remaining | *B. rapa* and *A. thaliana* CM Filtered | % Remaining | Mapped to R-o-18 and 19-26nt Small RNAs filtered | % Remaining |
| --- | --- | --- | --- | --- | --- | --- | --- | --- | --- | --- | --- | --- |
| WTR-o-18 | 17dpf endosperm | 1 | SRX7113618 | 14,782,390 | 6,319,548 | 42.75% | 6,319,238 | 99.9951% | 6,210,368 | 98.277% | 4,033,404 | 64.95% |
| WTR-o-18 | 17dpf endosperm | 2 | SRX7113619 | 23,412,918 | 9,108,385 | 38.90% | 9,107,867 | 99.9943% | 8,963,673 | 98.417% | 6,033,604 | 67.31% |
| WTR-o-18 | 17dpf endosperm | 3 | SRX7113620 | 14,579,403 | 5,827,576 | 39.97% | 5,827,156 | 99.9928% | 5,718,292 | 98.132% | 3,618,539 | 63.28% |
| WTR-o-18 | 17dpf embryo | 1 | SRX7113621 | 21,823,191 | 15,937,453 | 73.03% | 15,936,988 | 99.9971% | 15,445,456 | 96.916% | 10,619,292 | 68.75% |
| WTR-o-18 | 17dpf embryo | 2 | SRX7113622 | 26,629,773 | 18,404,570 | 69.11% | 18,404,128 | 99.9976% | 17,914,651 | 97.340% | 12,031,834 | 67.16% |
| WTR-o-18 | 17dpf embryo | 3 | SRX7113623 | 21,903,128 | 13,863,523 | 63.29% | 13,863,171 | 99.9975% | 13,459,146 | 97.086% | 9,174,737 | 68.17% |
| WTR-o-18 | Leaves | 1 | SRX7113689 | 27,441,990 | 10,767,807 | 39.24% | 10,767,489 | 99.9970% | 8,391,872 | 77.937% | 3,635,851 | 43.33% |
| WTR-o-18 | Leaves | 2 | SRX7113690 | 33,992,611 | 11,177,367 | 32.88% | 11,176,716 | 99.9942% | 9,002,798 | 80.550% | 4,033,874 | 44.81% |
| WTR-o-18 | Leaves | 3 | SRX7113691 | 29,436,807 | 9,381,536 | 31.87% | 9,381,375 | 99.9983% | 7,546,841 | 80.445% | 3,146,617 | 41.69% |
| WTR-o-18xR-o-18 | Embryo | 1 | SRX7113624 | 21,304,038 | 4,941,922 | 23.20% | 4,941,786 | 99.9972% | 4,804,648 | 97.225% | 3,098,803 | 64.50% |
| WTR-o-18xR-o-18 | Embryo | 2 | SRX7113625 | 16,203,619 | 4,132,008 | 25.50% | 4,131,841 | 99.9960% | 4,014,630 | 97.163% | 2,726,752 | 67.92% |
| WTR-o-18xR-o-18 | Embryo | 3 | SRX7113626 | 18,581,017 | 2,302,004 | 12.39% | 2,301,886 | 99.9949% | 2,199,864 | 95.568% | 1,446,836 | 65.77% |

**Supplemental Table 3.** Pearson correlation coefficients, r, for endosperm to embryo methylation.

|  | absolute<br>methylation (%) | fold change | difference in<br>methylation (%) |
| --- | --- | --- | --- |
| CG | 0.53 | -0.19 | -0.36 |
| CHG | 0.69 | -0.093 | -0.48 |
| CHH | 0.82 | 0.38 | 0.19 |

Correlation between endosperm methylation and mature embryo CHH methylation at CHH hyper-DMWs
|  | absolute<br>methylation (%) | fold change | difference in<br>methylation (%) |
| --- | --- | --- | --- |
| CG | 0.37 | 0.025 | -0.23 |
| CHG | 0.45 | 0.075 | -0.35 |
| CHH | 0.60 | 0.51 | 0.34 |

## References

1. Jones PA. Functions of DNA methylation: Islands, start sites, gene bodies and beyond [Internet]. Nature Reviews Genetics. Nature Publishing Group; 2012. p. 484–92. Available from: www.nature.com/reviews/genetics

2. Zhang H, Lang Z, Zhu JK. Dynamics and function of DNA methylation in plants [Internet]. Nature Reviews Molecular Cell Biology. Nature Publishing Group; 2018. p. 489–506. Available from: https://pubmed.ncbi.nlm.nih.gov/29784956/

3. Matzke MA, Mosher RA. RNA-directed DNA methylation: An epigenetic pathway of increasing complexity [Internet]. Nature Reviews Genetics. Nature Publishing Group; 2014. p. 394–408. Available from: http://dx.doi.org/10.1038/nrg3683

4. Zhong X, Hale CJ, Law JA, Johnson LM, Feng S, Tu A, et al. DDR complex facilitates global association of RNA polymerase v to promoters and evolutionarily young transposons. Nat Struct Mol Biol. Nat Struct Mol Biol; 2012;19:870–5.

5. Li Q, Gent JI, Zynda G, Song J, Makarevitch I, Hirsch CD, et al. RNA-directed DNA methylation enforces boundaries between heterochromatin and euchromatin in the maize genome. Proc Natl Acad Sci U S A. National Academy of Sciences; 2015;112:14728–33.

6. Blevins T, Podicheti R, Mishra V, Marasco M, Wang J, Rusch D, et al. Identification of pol IV and RDR2-dependent precursors of 24 nt siRNAs guiding de novo DNA methylation in arabidopsis. Elife [Internet]. eLife Sciences Publications Ltd; 2015;4. Available from: http://dx.doi.org/10.7554/eLife.09591

7. Zhai J, Bischof S, Wang H, Feng S, Lee TF, Teng C, et al. A one precursor one siRNA model for pol IV-dependent siRNA biogenesis. Cell. Cell Press; 2015;163:445–55.

8. Singh J, Mishra V, Wang F, Huang HY, Pikaard CS. Reaction Mechanisms of Pol IV, RDR2, and DCL3 Drive RNA Channeling in the siRNA-Directed DNA Methylation Pathway. Mol Cell. Cell Press; 2019;75:576–589.e5.

9. Cao X, Jacobsen SE. Role of the Arabidopsis DRM methyltransferases in de novo DNA methylation and gene silencing. Curr Biol. Cell Press; 2002;12:1138–44.

10. Wierzbicki AT, Ream TS, Haag JR, Pikaard CS. RNA polymerase v transcription guides ARGONAUTE4 to chromatin. Nat Genet. Nature Publishing Group; 2009;41:630–4.

11. Havecker ER, Wallbridge LM, Hardcastle TJ, Bush MS, Kelly KA, Dunn RM, et al. The arabidopsis RNA-directed DNA methylation argonautes functionally diverge based on their expression and interaction with target loci. Plant Cell. American Society of Plant Biologists; 2010;22:321–34.

12. Ye R, Wang W, Iki T, Liu C, Wu Y, Ishikawa M, et al. Cytoplasmic Assembly and Selective Nuclear Import of Arabidopsis ARGONAUTE4/siRNA Complexes. Mol Cell. Cell Press; 2012;46:859–70.

13. Greaves I, Groszmann M, Dennis ES, Peacock WJ. Trans-chromosomal methylation. Epigenetics. Taylor and Francis Inc.; 2012;7:800–5.

14. Hollick JB. Paramutation: A trans-homolog interaction affecting heritable gene regulation [Internet]. Current Opinion in Plant Biology. Elsevier Current Trends; 2012. p. 536–43. Available from: http://dx.doi.org/10.1016/j.pbi.2012.09.003

15. Aufsatz W, Mette MF, Van Der Winden J, Matzke AJM, Matzke M. RNA-directed DNA methylation in Arabidopsis. Proc Natl Acad Sci U S A. National Academy of Sciences; 2002;99:16499–506.

16. Molnar A, Melnyk CW, Bassett A, Hardcastle TJ, Dunn R, Baulcombe DC. Small silencing RNAs in plants are mobile and direct epigenetic modification in recipient cells. Science. American Association for the Advancement of Science; 2010;328:872–5.

17. Hollick JB. Paramutation and Development. Annu Rev Cell Dev Biol. Annual Reviews; 2010;26:557–79.

18. Gouil Q, Baulcombe DC. DNA Methylation Signatures of the Plant Chromomethyltransferases. Mittelsten Scheid O, editor. PLoS Genet. Public Library of Science; 2016;12:e1006526.

19. Grover JW, Kendall T, Baten A, Burgess D, Freeling M, King GJ, et al. Maternal components of RNA-directed DNA methylation are required for seed development in Brassica rapa. Plant J. Blackwell Publishing Ltd; 2018;94:575–82.

20. Chow HT, Chakraborty T, Mosher RA. RNA-directed DNA Methylation and sexual reproduction: expanding beyond the seed [Internet]. Current Opinion in Plant Biology. Elsevier Ltd; 2020. p. 11–7. Available from: http://dx.doi.org/10.1016/j.pbi.2019.11.006

21. Xu L, Yuan K, Yuan M, Meng X, Chen M, Wu J, et al. Regulation of Rice Tillering by RNA-Directed DNA Methylation at Miniature Inverted-Repeat Transposable Elements. Mol Plant. Cell Press; 2020;13:851–63.

22. Bouyer D, Kramdi A, Kassam M, Heese M, Schnittger A, Roudier F, et al. DNA methylation dynamics during early plant life. Genome Biol. BioMed Central Ltd.; 2017;18:179.

23. Kawakatsu T, Nery JR, Castanon R, Ecker JR. Dynamic DNA methylation reconfiguration during seed development and germination. Genome Biol. BioMed Central Ltd.; 2017;18:171.

24. Narsai R, Gouil Q, Secco D, Srivastava A, Karpievitch YV, Liew LC, et al. Extensive transcriptomic and epigenomic remodelling occurs during Arabidopsis thaliana germination. Genome Biol. BioMed Central Ltd.; 2017;18:172.

25. Lin JY, Le BH, Chen M, Henry KF, Hur J, Hsieh TF, et al. Similarity between soybean and Arabidopsis seed methylomes and loss of non-CG methylation does not affect seed development. Proc Natl Acad Sci U S A. National Academy of Sciences; 2017;114:E9730–9.

26. Rajkumar MS, Gupta K, Khemka NK, Garg R, Jain M. DNA methylation reprogramming during seed development and its functional relevance in seed size/weight determination in chickpea. Communications Biology. Nature Research; 2020;3:1–13.

27. Ji L, Mathioni SM, Johnson S, Tucker D, Bewick AJ, Kim KD, et al. Genome-wide reinforcement of DNA methylation occurs during somatic embryogenesis in soybean. Plant Cell. American Society of Plant Biologists; 2019;31:2315–31.

28. Bauer MJ, Fischer RL. Genome demethylation and imprinting in the endosperm. Curr Opin Plant Biol. 2011;14:162–7.

29. Lafon-Placette C, Köhler C. Embryo and endosperm, partners in seed development. Curr Opin Plant Biol. 2014;17:64–9.

30. Mosher RA, Melnyk CW. siRNAs and DNA methylation: seedy epigenetics. Trends Plant Sci. 2010;15:204–10.

31. Grover JW, Burgess D, Kendall T, Baten A, Pokhrel S, King GJ, et al. Abundant expression of maternal siRNAs is a conserved feature of seed development. Proc Natl Acad Sci U S A. NLM (Medline); 2020;117:15305–15.

32. Gehring M, Bubb KL, Henikoff S. Extensive demethylation of repetitive elements during seed development underlies gene imprinting. Science. American Association for the Advancement of Science; 2009;324:1447–51.

33. Hsieh TF, Ibarra CA, Silva P, Zemach A, Eshed-Williams L, Fischer RL, et al. Genome-wide demethylation of Arabidopsis endosperm. Science. 2009;324:1451–4.

34. Zemach A, Kim MY, Silva P, Rodrigues JA, Dotson B, Brooks MD, et al. Local DNA hypomethylation activates genes in rice endosperm. Proc Natl Acad Sci U S A. National Academy of Sciences; 2010;107:18729–34.

35. Zemach A, Kim MY, Hsieh PH, Coleman-Derr D, Eshed-Williams L, Thao K, et al. The arabidopsis nucleosome remodeler DDM1 allows DNA methyltransferases to access H1-containing heterochromatin. Cell. 2013;153:193–205.

36. Wang Z, Baulcombe DC. Transposon age and non-CG methylation. Nat Commun. Nature Research; 2020;11:1–9.

37. Nowack MK, Ungru A, Bjerkan KN, Grini PE, Schnittger A. Reproductive cross-talk: Seed development in flowering plants [Internet]. Biochemical Society Transactions. Portland Press; 2010. p. 604–12. Available from: https://portlandpress.com/biochemsoctrans/article-pdf/38/2/604/517309/bst0380604.pdf

38. Figueiredo DD, Köhler C. Signalling events regulating seed coat development [Internet]. Biochemical Society Transactions. Portland Press Ltd; 2014. p. 358–63. Available from: https://portlandpress.com/biochemsoctrans/article-pdf/42/2/358/487379/bst0420358.pdf

39. Iwasaki YW, Siomi MC, Siomi H. PIWI-interacting RNA: Its biogenesis and functions. Annu Rev Biochem. Annual Reviews Inc.; 2015;84:405–33.

40. Martínez G, Panda K, Köhler C, Slotkin RK. Silencing in sperm cells is directed by RNA movement from the surrounding nurse cell. Nat Plants. 2016;2:16030.

41. Martinez G, Wolff P, Wang Z, Moreno-Romero J, Santos-González J, Conze LL, et al. Paternal easiRNAs regulate parental genome dosage in Arabidopsis. Nat Genet. 2018;50:193–8.

42. Papareddy RK, Páldi K, Paulraj S, Kao P, Nodine MD. Chromatin Regulates Bipartite-Classified Small RNA Expression to Maintain Epigenome Homeostasis in Arabidopsis [Internet]. 2020 [cited 2020 Sep 2]. p. 2020.05.04.076885. Available from: https://www.biorxiv.org/content/10.1101/2020.05.04.076885v1.abstract

43. Urich MA, Nery JR, Lister R, Schmitz RJ, Ecker JR. MethylC-seq library preparation for base-resolution whole-genome bisulfite sequencing. Nat Protoc. Nature Publishing Group; 2015;10:475–83.

44. Babraham Bioinformatics – FastQC A Quality Control tool for High Throughput Sequence Data [Internet]. [cited 2020 Sep 2]. Available from: https://www.bioinformatics.babraham.ac.uk/projects/fastqc/

45. Andrews S. FastQC [Internet]. Github; [cited 2020 Sep 2]. Available from: https://github.com/s-andrews/FastQC

46. Babraham Bioinformatics – Trim Galore! [Internet]. [cited 2020 Sep 2]. Available from: https://www.bioinformatics.babraham.ac.uk/projects/trim_galore/

47. Pedersen BS, Eyring K, De S, Yang IV, Schwartz DA. Fast and accurate alignment of long bisulfite-seq reads. 2014; Available from: http://arxiv.org/abs/1401.1129

48. Picard Tools – By Broad Institute [Internet]. Available from: http://broadinstitute.github.io/picard/

49. Li H, Handsaker B, Wysoker A, Fennell T, Ruan J, Homer N, et al. The Sequence Alignment/Map format and SAMtools. Bioinformatics. Oxford Academic; 2009;25:2078–9.

50. Pedersen BS, Quinlan AR. Mosdepth: Quick coverage calculation for genomes and exomes. Bioinformatics. Oxford University Press; 2018;34:867–8.

51. dpryan79/MethylDackel: A (mostly) universal methylation extractor for BS-seq experiments. [Internet]. Available from: https://github.com/dpryan79/methyldackel

52. Quinlan AR. BEDTools: The Swiss-Army tool for genome feature analysis. Curr Protoc Bioinformatics. John Wiley and Sons Inc.; 2014;2014:11.12.1–11.12.34.

53. Quinlan AR, Hall IM. BEDTools: A flexible suite of utilities for comparing genomic features. Bioinformatics. Oxford Academic; 2010;26:841–2.

54. boseHere/sRNA_snakemake_workflow: A snakemake workflow for an sRNA-seq data analysis pipeline developed by the Mosher Lab [Internet]. Available from: https://github.com/boseHere/sRNA_snakemake_workflow

55. Axtell MJ. ShortStack: Comprehensive annotation and quantification of small RNA genes. RNA. Cold Spring Harbor Laboratory Press; 2013;19:740–51.

56. Johnson NR, Yeoh JM, Coruh C, Axtell MJ. Improved placement of multi-mapping small RNAs. G3: Genes, Genomes, Genetics. Genetics Society of America; 2016;6:2103–11.

